# Methylglyoxal Induces Endothelial Dysfunction via Stunning-like Phenotype

**DOI:** 10.1101/2021.11.18.469085

**Authors:** Thomas Fleming, Bastian von Nettelbladt, Jakob Morgenstern, Marta Campos, Maxime Le Marois, Maria Bartosova, Ingrid Hausser, Constantin Schwab, Andreas Fischer, Peter P. Nawroth, Julia Szendroedi, Stefan Herzig

## Abstract

Elevated levels of methylglyoxal (MG) and its associated post-translational modifications have been reported to be associated with progression and development of numerous pathological conditions. Despite such extensive evidence, it still remains unclear what the specific effects of MG are other than that induction of cytotoxicity. Here we evaluated the effects of MG in cardiac endothelial cells *in vitro*. We found that MG leads to a non-proliferative state and endothelial dysfunction, which is reversible as MG-H1, a major post-translational modification induced by MG, is turned over by lysosomal degradation. MG-induced cellular stunning/paralysis describes a new hallmark for cellular dysfunction which could lead to alterations in tissue homeostasis as well as cell-to-cell interactions, thereby contributing to the pathogenesis of diseases.

## Introduction

Methylglyoxal (MG), a by-product formed from the non-enzymatic degradation of triosephosphate pool of metabolites in glycolysis, belongs to class of endogenous stressors referred to as the reactive carbonyl species (RCS), and acts as a potent glycating agent to form advanced glycation endproducts (AGEs) (Thornalley, 2008; Rabbani and Thornalley, 2012).

Studies have shown that modification of human serum albumin, vascular basement membrane type IV collagen and apolipoprotein B100 of low-density lipoprotein by MG is leading to the formation of the arginine-derived hydroimidazolone AGE, MG-H1, can cause structural distortions, loss of side chain charge, and functional impairment (Ahmed et al., 2005; Dobler et al., 2006; Rabbani et al., 2011). It has also been reported that free MG-H1, the adduct residue released from modified proteins by cellular proteolysis, can be detected in plasma, urine, cerebrospinal fluid, synovial fluid, as well as peritoneal dialysis and elevated levels have been associated with advanced end-stage renal disease, cirrhosis, Alzheimer’s disease, Parkinson’s disease and aging (Rabbani and Thornalley, 2012). Modification of the DNA by MG has been shown to be associated with DNA double-strand breaks, whilst increased excretion of the modified nucleoside has been reported in the plasma and urine of patients with diabetic nephropathy (Thornalley et al., 2010a; Li et al., 2006; Waris et al., 2015).

Despite extensive *in vivo* evidence, it still remains unclear what the *in vitro* effects of MG are other than the induction of cytotoxicity (Ayoub et al., 1993; Belanger et al., 2011; Chu et al., 2016; de Oliveira et al., 2019; di Emidio et al., 2019; Galligan et al.; Paramita and Wisnubroto, 2018; Prantner et al., 2021; Roy et al., 2018; Sachdeva et al.; Suh et al., 2018a; b). Due to the capacity of MG to modify arginine, it would be expected that such effects would be damaging, as arginine residues have a high probability of being located at functional relevant sites such as those involved in protein-protein interactions, enzyme-substrate interactions, as well as protein-DNA interactions. It has been shown that whilst detection and identification of MG-H1 modified proteins is possible, the extent of modification, even under physiological relevant increases of MG, is very low (Rabbani and Thornalley, 2014a; Irshad et al., 2019).

Here we report that when intracellular MG is transiently increased 12-fold, relative to the basal level, there is an equivalent increase in MG-H1, localized to the nucleus. This leads to a non-proliferative and unresponsive state that was not associated with cell death, quiescence or senescence, but the induction of endothelial dysfunction. The inability of the cell to respond to extracellular stimuli would suggest MG effects chromatin dynamics, leading to cell cycle arrest and the loss of proliferative capacity. This cellular phenotype was found to be reversible as MG-H1 was turned over by lysosomal degradation. MG-induced cellular stunning/paralysis therefore represents a novel type of cellular dysfunction which could lead to alterations in tissue homeostasis as well as cell-to-cell interactions, thereby contributing to the pathogenesis of diseases, particularly those conditions in which MG can accumulate.

## Results

### Cellular Metabolism of Exogenous Methylglyoxal

The basal levels of MG in murine cardiac endothelial cells (MCECs) was 42.6 ± 4.7μM, equivalent to 32.0 ± 3.5 pmol/10^6^ cells. The uptake of deuterium labelled MG ([d_4_]-MG) was rapid; one-hour post-stimulation, the intracellular concentration of [d4]-MG correlated positively with the exogenous concentrations, in a proportional manner (**Figure 1A**). At 500μM MG, the intracellular level was increased 12-fold relative to basal levels. After one-hour, the MG was rapidly removed from the intracellular compartment, following first-order kinetics. After 12hrs, the intracellular concentration of [d4]-MG at all of the stated exogenous concentrations had decreased below the basal level of MG.

**Figure 1.**
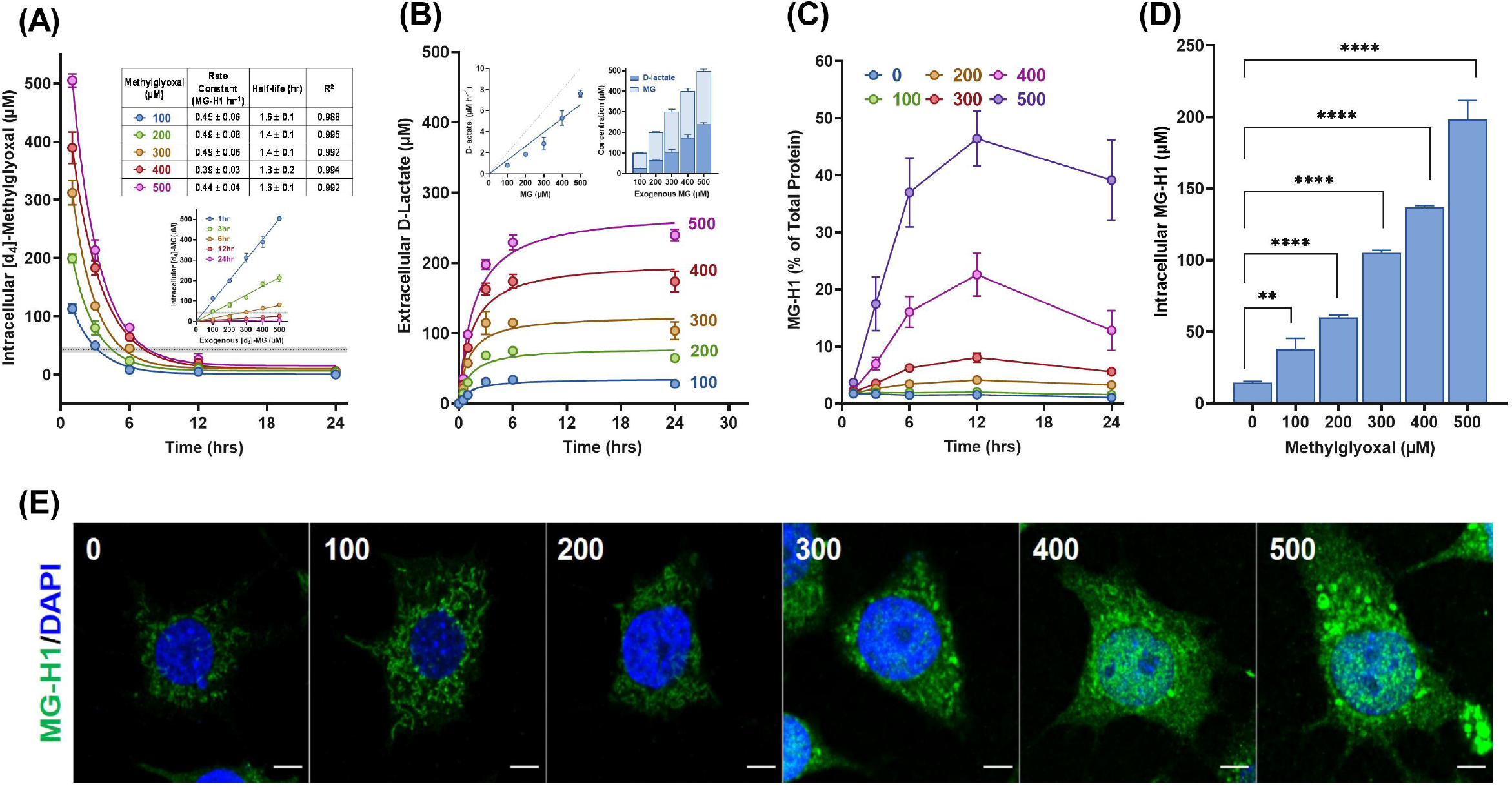
Cellular Metabolism of Exogenous Methylglyoxal. Intracellular [d4]-MG concentrations following stimulation with [d4]-MG (**A**) were measured as described in the material and methods section. Rate constants and half-life given in the inset table were calculated by fitting of the data to a first-order decay using GraphPad Prism^®^ software. The inset graph shows the linear relationship between the exogenous and intracellular concentration. The shaded line in each plot represents the mean concentration of MG measured at each time point in the MCECs. Data represent mean ± SD (N=3). (**B**) The relationship between exogenous MG (0-500μM) and the production of extracellular D-lactate in vitro. The data has been normalized to the levels of D-lactate measured in the unstimulated MCECs. The rate of D-lactate formation, shown in the *left* inset, was calculated by linear regression of the data from 0-3hrs using GraphPad Prism^®^ software. The dashed line indicates a directly proportional relationship. Theoretical amount of MG which would remain in the intracellular compartment following the conversion of the exogenous MG to D-lactate, shown in the *right* inset. Data represent mean ± SD (N=3). (**C**) Formation of MG-H1 in MCEC stimulated with MG. MCECs were seeded into 96-well plates at a density of 25000 cells per well in 200μl of medium containing 0.1% FCS. After 16hrs, the MCECs were stimulated with MG (0-500μM) and at consecutive interval, the plates/cells fixated and stained for MG-H1 as described in material and methods section. Data is expressed as % of the total protein and represents mean ± SD (N=8). (**D**) Quantification of intracellular MG-H1 by LC-MS/MS analysis. Data represent mean ± SD (N=3); ****P ≤ 0.0001, **P ≤ 0.01 vs. unstimulated MCECs. (**E**) Subcellular localization of MG-H1 in MCECs stimulated with MG (0-500μM) for 6hrs. MCECs were seeded onto gelatin coated 12mm glass cover slips in 24-well. After 16hrs, the media was exchanged to contain the specified concentrations MG and incubated for 6hrs. MCECs were then fixated, permeabilized and blocked, as described in the materials and methods section and then stained for MG-H1 (Green) and nuclei (Blue; Dapi). Images were acquired on a Zeiss LSM780 confocal microscope with a Plan-Apochromat 40x/1.3 Oil objective. Representative images are shown in the *upper* panel in which the scale bar represents 10μm.

The rate of D-lactate production correlated significantly with the MG levels but was not proportional (**Figure 1B**). Assuming that the conversion of MG to D-lactate by the glyoxalase system is stoichiometrically equivalent, then the lack of proportional detoxification by the glyoxalase system means that only 37% of MG is converted to D-lactate, the remainder of which is available to react with components of the intracellular compartments. In this respect, it was shown that the formation of MG-H1 was hyperbolic, with the start point of the plateau occurring at six-hours post-stimulation, regardless of the MG concentration, and remained relatively stable until 24hrs post-stimulation (**Figure 1C**). LC-MS/MS analysis showed that at six-hours, the amount of MG-H1 correlated positively with, and proportional to, the exogenous concentrations of MG (**Figure 1D**). There were no changes in any of other markers of glycation, oxidation or nitration. At 500μM MG, the amount of total MG-H1 had significantly increased by 13-fold consistent with the increase in the intracellular concentration of MG at one-hour. Immunofluorescence analysis showed that at this concentration, the accumulation of MG-H1 within the intracellular compartment was within the nucleus (**Figure 1E**).

### Exogenous Methylglyoxal Induces Loss of Proliferation Capacity

Stimulation of MCECs with MG leads to a dose-dependent reduction in cell viability after 24hrs (**Figure 2A**). Interestingly, the release of lactate dehydrogenase (LDH), a marker of membrane integrity and in turn cytotoxicity, did show a dose-dependent increase, however, the EC_50_ was 1.5-fold higher compared to the viability readouts. Furthermore, the magnitude of this increase was small as compared to other markers, suggesting that MG does result in the loss of membrane integrity and the induction of cytotoxicity is not an explanation for the changes observed in other markers of viability.

MG was subsequently found to induce cell death at concentrations >900μM after 24hrs (**Figure 2B** & **2C**) and was associated with the cleavage of the pro-apoptotic mediators, Caspase-3 and PARP, as well as the induction of DNA damage response pathway at concentrations >600μM (**Figure 2D**, *Upper panel*). Consistently, the proliferative capacities of the MCECs at >600μM showed an immediate reduction; 6hrs post-stimulation the normalized cell index had reduced to zero and did not increase over the remaining monitoring period (**Figure 2E**). At concentrations <600μM, MG induced a dose-dependent decrease in the growth rate constant, suggesting that a major physiological effect of MG is the loss of cellular viability through a decreased proliferative capacity. Consistently, MG was found to induce a dose-dependent decrease in the synthesis of DNA, RNA and protein (**Figure 2F**).

**Figure 2.**
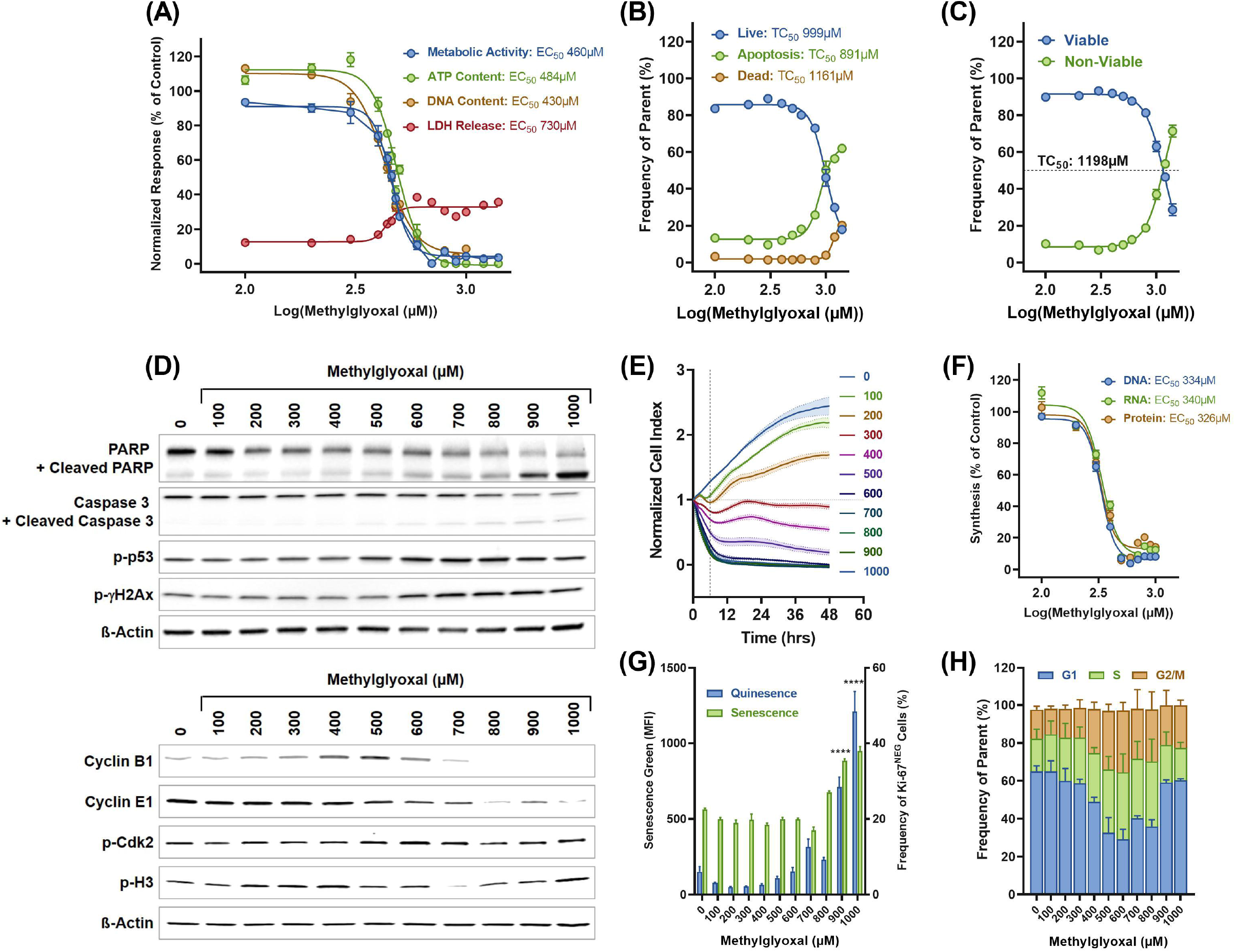
Methylglyoxal Induces Loss of Proliferative Capacity. The effect of MG on cell viability markers after 24hrs (**A**). Data represents mean ± SEM (*N* = 16-24) and EC_50_ curves generated using GraphPad Prism^®^ software. The cytotoxic effects of MG measured by flow cytometry for Annexin V (**B**) and live/dead discrimination (**C**). Data represents mean ± SD (*N* = 4) and EC_50_ curves generated using GraphPad Prism^®^ software. (**D**) Activation of the apoptosis, damage response pathway and cell cycle arrest were assessed by western blot of total extracts prepared from MCECs stimulated with MG for 24hrs, as described in the material and methods section, using the specified protein markers. ß-Actin was used as a loading control. (**E**) Proliferation analysis of MCECs stimulated with MG was measured using the xCELLigence RTCA platform, as described in the material and methods. The cell index was internally normalized to the cell index value measured prior to MG treatment. Data represents mean ± SD (N = 8 per conc.). (**F**) The inhibition of DNA, RNA and protein synthesis in MCECs stimulated with MG after 24hrs was measured as described in the material and methods. Data represents mean ± SD (N = 8) and EC_50_ curves generated using GraphPad Prism^®^ software. **(G)** Quiescence and Senescence in MCECs stimulated with MG (0-1000μM) was measured in the viable cell population, 24hrs post-stimulation, by flow cytometry analysis, as described in the materials and methods section. Data represents mean ± SD (N = 4); ****P ≤ 0.0001 vs. unstimulated MCECs. (**H**) Cell cycle analysis in MCECs stimulated with MG was measured 24hrs post-stimulation, by flow cytometric analysis using propidium iodide (PI), as described in the materials and methods section. Data represents mean ± SD (N = 4).

The loss of proliferative capacity was not associated with the induction of quiescence or senescence (**Figure 2G**). However, cell cycle analysis showed that the frequency of G1 cells in MCECs stimulated with 500μM of MG had significantly decreased by about 2-fold as compared to the untreated MCECs (**Figure 2H**). The loss of G1 cells was paralleled by a significant 2-fold increase in the frequency of S and G2/M cells, suggesting that MG induces an arrest at the S/G2-M phase of the cell cycle. Consistently, western blot analysis showed that after 24hrs post-treatment with MG, there was a reduction in the levels of Cyclin B and E, whilst phosphorylation of cdk2 and Histone H3 were also enhanced (**Figure 2D**,*lower panel*).

### Methylglyoxal Induces Endothelial Dysfunction

MCECs which have been stimulated with MG did not show any significant changes in morphology (**Figure 3A**), however, there was a significant decrease in the surface expression of CD106/VCAM-1, and significant increases in CD146/MUC18, CD201/EPCR and CD62E/E-selectin (**Figure 3B**). All of these changes, observed at 500μM MG, could be prevented by pre-incubation with the MG scavenger, aminoguanidine (AG), thereby confirming that these effects are as a direct result of MG. The MG stimulated MCECs also exhibited an increase in adhesiveness, not only with respect to leukocytes (**Figure 3C**) but also with respect to their adhesion to gelatine coated plastic (**Figure 3D**).

**Figure 3.**
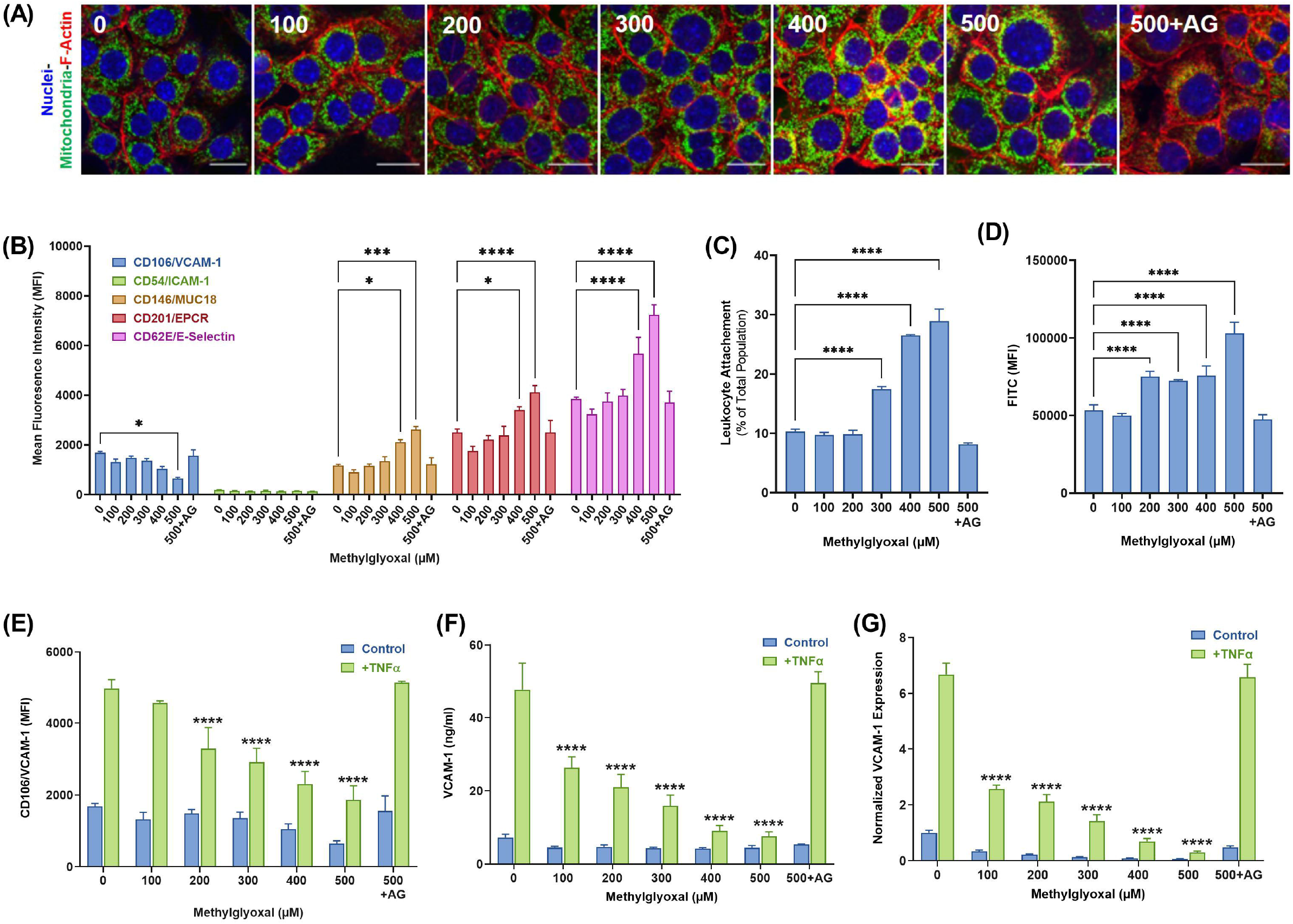
Methylglyoxal Induces Endothelial Dysfunction. Morphological analysis of MCECs stimulated with MG (0-500μM) of 24hrs by immunofluorescence (**A**). MCECs were seeded onto gelatin coated 12mm glass cover slips in 24-well. After 16hrs, the media was exchanged to contain the specified concentrations MG and incubated for 24hrs. MCECs were then fixated, permeabilized and blocked, as described in the materials and methods section and then stained for mitochondria (Green; Complex I), F-actin (Red; Phalloidin-iFluor 647) and nuclei (Blue; Dapi). Images were acquired on Zeiss LSM780 microscope with oil immersion at x40 magnification. Scale bar represents 20μm. (**B**) The activation status of MCECs stimulated with MG (0-500μM) was measured in the viable cell population, 24hrs post-stimulation, by flow cytometry analysis, as described in the material and methods section. Data represents mean ± SD (N = 4); ****P ≤ 0.0001, ***P ≤ 0.001, *P ≤ 0.05 vs. unstimulated MCECs. (**C**) Leukocyte adhesion of MCECs stimulated with MG (0-500μM) was measured, 24hrs post-stimulation, by flow cytometry analysis, as described in the materials and methods section. Data represents mean ± SD (N = 4); ****P ≤ 0.0001 vs. unstimulated MCECs. (**D**) General adhesion of CSFE-labelled MCECs stimulated with MG (0-500μM) to gelatin coated surface was measured, 24hrs post-stimulation, using fluorescence microplate reader, as described in the materials and methods section. Data represents mean ± SD (N = 4); ****P ≤ 0.0001 vs. unstimulated MCECs. (**E**) TNFα responsiveness of MCECs stimulated with MG (0-500μM). 24hrs post-stimulation, the media was exchanged to contain TNFα (10ng/ml). After 16hrs, the cells were harvested and analyzed for the surface expression of CD106/VCAM-1 by flow cytometry, as described on the material and methods. The media was analyzed for shed CD106/VCAM-1by ELISA (**F**) and the cells were also analyzed for mRNA expression of CD106/VCAM-1, by qPCR (**G**). Data represents mean ± SD (N = 4); ****P ≤ 0.0001 vs. unstimulated MCECs stimulated with TNFα.

The responsiveness of the MG stimulated MCECs to TNFα was also found to be impaired (**Figure 3E**). Whilst the expression of CD106/VCAM-1 was significantly increased following stimulation with TNFα, the magnitude of the increase in the MG stimulated MCECs was significantly decreased. This deficiency could not be explained by increased shedding of CD106/VCAM-1 from the cell surface into the media (**Figure 3F**); for CD106/VCAM-1 the levels in the media reflected the changes observed for the surface expression. However, the mRNA expression for CD106/VCAM-1 at baseline and following TNFα treatment did reflect the changes observed in the surface expression (**Figure 3G**).

Other key functions of endothelial cells such as the wound healing, angiogenesis and barrier function were also assessed (**Supplementary Figure 1**). The rate of wound closure was significantly reduced in the MCECs which has been stimulated with concentrations >300μM MG. Consistent with these results, the angiogenetic potential was also significantly reduced in the MCECs at the same concentration range. The deficits in wound healing and tube formation observed at 500μM MG could be prevented by pre-incubation with AG. MG stimulation was also found to impair barrier function by non-significantly reducing transendothelial resistance, it did not affect permeability to large molecular compounds.

### Effects of Methylglyoxal Stimulation are Recoverable

The effects of MG, specifically the loss of proliferation, were found to be reversible. It was found that 48hrs after stimulation with 500μM MG, MCECs continued to show a significant 8-fold reduction in the growth rate constant and a concurrent increase in the doubling time, as compared to unstimulated MCECs (**Figure 4A**). However, after 48hrs, the proliferation capacity began to normalize, such that by the end of the monitoring period, there was no-significant difference with respect to the growth rate constant and/or doubling time (**Figure 4B**). Furthermore, the responsiveness of the MG stimulated MCECs to TNFα had also recovered (**Figure 4C**).

**Figure 4.**
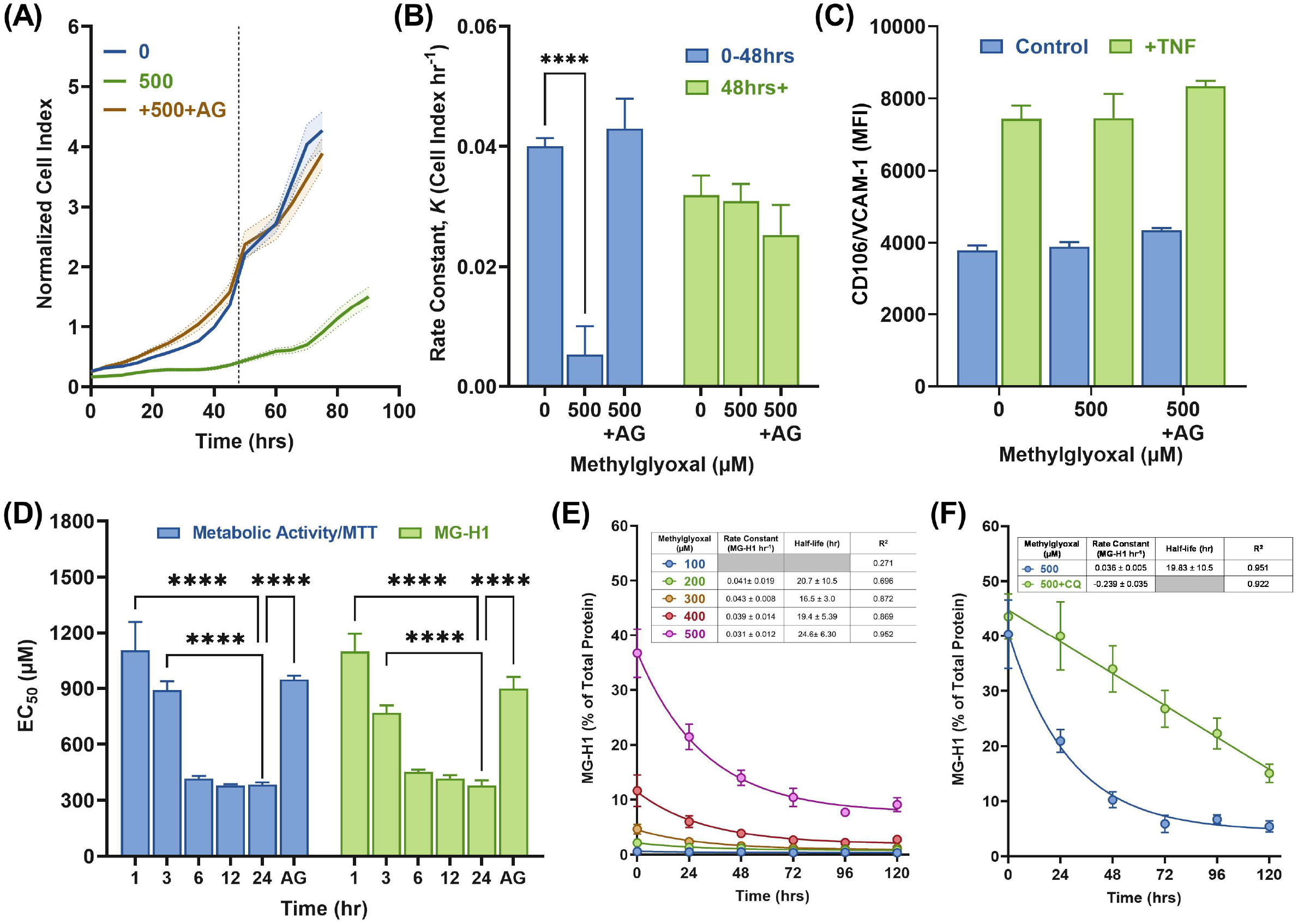
Cellular Effects of Methylglyoxal Stimulation are Reversible. The proliferative capacity of MCECs which had been stimulated with MG for 24hrs was measured using the xCELLigence RTCA platform as described in the material and methods section (**A**). The cell index was internally normalized to the cell index value measured at 12hrs post-seeding. Data represents mean ± SD (N = 4). The dashed line indicates the 48hr time-point. (**B**) Growth constant (k) calculated from the proliferation profile by fitting to an exponential (Malthusian) growth curve using GraphPad Prism^®^ software. Data represents mean ± SD (N = 4); ****P ≤ 0.0001 vs. unstimulated MCECs. (**C**) TNFα responsiveness of MG stimulated MCECs, 72hrs post-stimulation, was assessed as described in the material and methods. Data represents mean ± SD (N = 3). The effect of media exchange on total metabolic activity, as measured by MTT and MG-H1 formation (**D**). Data represent mean ± SD (N=8 per time point); ****P ≤ 0.0001 vs. 24hr time point. (**E**) The intracellular turnover of MG-H1 in MCECs stimulated with MG for 24hrs. Data is expressed as % of the total protein and represents mean ± SD (N=8). Rate constants and half-life given in the inset table were calculated by fitting of the data to a first-order decay using GraphPad Prism^®^ software. (**F**) The effect of lysosomal inhibition on the degradation of MG-H1. Data is expressed as % of the total protein and represents mean ± SD (N=8). Rate constants and half-life given in the inset table were calculated by fitting of the data to either a first-order or zero-order decay using GraphPad Prism^®^ software.

In order to establish whether this recoverability was associated to the extent of MG-H1 modification, the relationship between MG-H1 and the cellular effects induced by MG would first need to be demonstrated. It was shown that exchanging MG containing media after one and three-hours significantly prevented the reduction in total metabolic activity (**Figure 4D**), DNA content and the inhibition of macromolecule synthesis (**Supplementary Figure 2**). However, exchanging the media after six and twelve-hours had no significant effect. With respect to MG-H1, a similar trend was also observed, in that the removal of the MG containing media at the early time points prevented the level of MG-H1 observed at 24hrs, equivalent to the effect of pre-incubation with AG (**Figure 4D**). These results show that the cellular effects observed with MG are dependent upon the formation of MG-H1.

The turnover of MG-H1 was also measured in MCECs which had been stimulated with MG for 24hrs. It was found that the degradation of MG-H1 followed first order kinetics (**Figure 4E**). After 120hrs, the MG-H1 content of the stimulated cells had decreased, however at concentrations >300μM, the MG-H1 content still remained significantly elevated as compared to the unstimulated MCECs. The turnover of MG-H1 was shown not to be dependent on the proteasome, as the proteasomal inhibitor, MG132(Lee and Goldberg, 1998), had no significant effect. It was subsequently shown that the activity of the proteasome, as well as the proteases which constitute its activity were inhibited in MG stimulated MCECs. Furthermore, electron microscopy analysis of cells showed no ultrastructural evidence of protein aggregation as indicated by the absence of electron dense areas in the MG stimulated MCECs, as compared to MCECs which had been treated with MG132 alone (**Supplementary Figure 2**). Subsequent screening with different inhibitors of the protein degradation pathway identified the lysosomal inhibitor, chloroquine (CQ) (Bonam et al., 2019), as being the only effective means of reducing the degradation of the MG-H1 (**Figure 4F**).

## Discussion

Elevated levels of MG and its associated post-translational modifications have been reported to be associated with progression and development of numerous pathological conditions (Rabbani and Thornalley, 2012). It still remains unclear, despite extensive studies, what the cellular effects of MG are and how they are induced within the intracellular environment. In this study, the effects of MG on MCECs were studied. It was found that the uptake and elimination of MG from the intracellular compartment was rapid, reaching an equal molarity to the exogenous concentration one-hour post-stimulation. A proportion of the intracellular MG was subsequently removed through detoxification by the glyoxalase system. However, the majority of MG resulted in protein modification with a maximum MG-H1 formation occurring six-hours post-stimulation regardless of the intracellular concentrations at one-hour. The lag time between these two events is likely due to the kinetics of MG diffusing into the intracellular compartment and reacting with accessible arginine residues (Lo et al., 1994). After six-hours, the level of modification remained relatively stable up to 24hrs post-stimulation. At this time, many cellular effects could be observed that can be broadly classified as *endothelial dysfunction,* an initiating factor for pathogenesis of many cardiovascular diseases (Avogaro et al., 2011). Many of these effects were evident at 500μM, when the intracellular concentration of MG was 12-fold higher than the basal levels. Under this condition, there was an equivalent increase in MG-H1, localized to the nucleus. Preventing the increase in MG, and in turn MG-H1, also prevented the cellular effects described. The relative increases observed at 500μM can therefore be viewed as the intracellular threshold that needs to be achieved in order for the MG/MG-H1 to induce endothelial dysfunction. Increases above this threshold were associated with the induction of DNA damage and cytotoxicity, indicating the narrowness between biological and toxicity effects of MG/MG-H1.

A key feature resulting from MG stimulation was the inability of the cell to effectively respond to a proinflammatory stimuli. This manifested itself primarily at the transcriptional level as the changes in transcribed mRNA reflected, to some degree, the extent of translated protein. This would suggest that MG/MG-H1 affects chromatin dynamics. The localization of MG-H1 to the nucleus would be consistent with this conclusion, and has been reported previously (Wang et al., 2015). It has been shown that modification of the DNA by MG can lead to DNA strand breaks (Thornalley et al., 2010b). In this study, the induction of DNA damage was associated with the use of concentrations >600μM. Although no measurement of MG-derived DNA modifications was made, it would be expected that such modifications would occur at these concentrations leading to the activation of the DNA damage response and ultimately cytotoxicity. It has also been shown that modification of histones by glycation interfere with the epigenetic signature of a cell, thereby changing the regulatory functions in the processing of genetic information, independent of any direct effects that MG might have on the DNA (Rahmanpour and Bathaie, 2011; Zheng et al., 2019; Maksimovic and David, 2021; Talasz et al., 2002; Gugliucci, 1994; Galligan et al., 2018). Modification of the nuclear proteins, rather than the DNA, could therefore be the driving force behind the effects induced by MG. A blockage in transcription and translation, resulting from a change in chromatin dynamics, would be consistent with the observed inhibition in the synthesis of the macromolecules, which occurred at an EC_50_ that proceeded the cell arrest, the loss in proliferative capacity and cell viability markers. If this were the case, then such an effect could be considered as universal as the flow of genetic information is a process that occurs in all nucleated cells. However, it would be expected that the resulting effects would be cell specific. Further studies, using proteomics, would be required to identify the nuclear proteins which have been modified by MG-H1, as means of further identifying the mechanism of action.

The intracellular turnover of MG-H1, leading to the recovery from endothelial dysfunction were not dependent upon the proteasome, but lysosomal degradation. It has previously been proposed that oxidized proteins, including those which have been modified by AGEs, form aggregate like structures which are subsequently degraded by the proteasome (Jung and Grune, 2013; Jung et al., 2014). In this study it was shown that both the activity of the proteasome and the proteases which constitute such activity were inhibited by MG. This is consistent with a previous study which reported the proteasome as target for glycation, leading to the loss of activity (Queisser et al., 2010). The reduction in activity observed in this study could not be directly related to modification by MG-H1. There was also no evidence for the formation of morphologically observable aggregates. The contribution of the lysosomal pathway which includes the cathepsins, a superfamily of proteases known for their role in the degradation of AGE-modified proteins in vivo (Brings et al., 2017; Grimm et al., 2012), would suggest that the turn-over of MG-H1 proteins is more related to the loss in protein functionality, rather than the capacity to form large insoluble aggregates. Further studies, are required, however, to determine whether this is the case, particularly with respect to DJ1, which has been proposed as a MG-H1 specific deglycosylase (Richarme et al., 2015).

Cells can respond to stress in a variety of ways, ranging from the activation of the pathways which promote survival to eliciting programmed cells death to remove the damaged cells. These processes reflect the two outcomes of *allostasis,* or the process of maintaining homeostasis (Galluzzi et al., 2018; Fulda et al., 2010). In-between these two outcomes, there are different non-proliferative states in which a cell can enter to either reduce an/or repair the damage resulting from a stressor. Two examples are quiescence, which occurs due to lack of nutrition and growth factors, and senescence, which takes place due to aging and excessive DNA damage (Terzi et al., 2016). In both states, proliferation is halted through cell cycle arrest initiated by intrinsic cues resulting from the activity of mechanistic target of rapamycin (mTOR) and tumour-suppressor protein, p53. In quiescence, the cell cycle arrest occurs at G0 and is reversible whereas senescence can occur at any of the check points within the cell cycle, resulting ultimately in cell death (Terzi et al., 2016). With respect to MG, neither quiescence or senescence were induced at the concentrations associated with the cellular effects observed. However, at the same concentrations, the cells did enter S/G2-M arrest and lost proliferative capacity. This state was also reversible because as MG-H1 was turned over, the proliferative capacity was regained, suggesting that this cell state is more equivalent to the phenotype of quiescence. For this reason, the cellular effects of MG could be defined as *cellular stunning*, which refers to the period of time over which the capability of the cell to self-stabilize in response to MG – the so-called *homeostatic capacity*-is first lost through the formation of MG-H1 and then regained, through its turnover. A similar phenotype was described in CD8+ T cells, resulting from the transfer of MG from myeloid-derived suppressor cells via direct cell contact (Baumann et al., 2020). This led to the loss of effector function in the CD8+ T cells. Due to the temporary nature of the loss in function, the authors referred to the effect as *paralysis* rather than stunning. Furthermore, the loss and regain of function in CD8+T cells was associated with the levels of L-arginine, rather than extent of modification by MG-derived AGEs and/or the localization of such modifications to the nucleus.

The findings from this current study help to demonstrate that depending upon the concentration, as well as the cellular context, the effects from an acute exposure to MG can be more nuanced and impactful than the extreme response of cell death. It remains to be determined, however, whether MG-stunning/paralysis occurs *in vivo*. The quantification of MG and/or protein-bound MG-H1 within a given tissue would only provide information as to whether the steady levels were increased. As has been shown in this study, the intracellular MG levels only need to be increased transiently to induce the stunning/paralysis phenotype. The frequency of nuclei MG-H1 within tissue could provide an indirect mean of identification. Preliminary analysis would suggest that in normal tissue, the frequency of nuclear MG-H1 is 9.4 ± 7.4% which can increase to 30-60% under disease conditions (**Supplementary Figure 3**). This increase could be indicative of a causative effect or it could merely represent a consequence resulting from an already established pathological condition. If it is causative, then the accumulation of MG-stunned/paralyzed cells could lead to the alterations to tissue homeostasis as well as cell-to-cell interactions, thereby contributing to the pathogenesis of diseases, particularly those conditions in which MG can accumulate.

## Material & Methods

### Chemicals

Methylglyoxal solution (40% w/v) was purchased from Sigma. [d4]-Methylglyoxal was synthesized, as previously described from [d6]-acetone (Clelland and Thornalley, 1990). The concentrations of the stock MG solutions were determined by LC-MS/MS and validated, along with the purity by 13C- and 1H-NMR using 298 K Bruker Advance II NMR spectrometer, prior to use in any cell culture experiments. The purity of the stock solution was 60-65% with the major contaminants being acetate (5.1%) methanol (0.4%), and formate (0.3%). A working solution (≥100mM) was prepared by dilution into distilled water, which was subsequently diluted into the media.

### Cell Culture

Murine cardiac endothelial cells (MCECs) were purchased from Biozol/CELLutions Biosystems Inc (Catalogue No. CLU510). Cells were grown on 0.5% gelatin-coated surface in DMEM with 5% FCS, 1 g/l glucose, 1% penicillin/streptomycin, 1% amphotericin B, 10mM HEPES at 37°C. Cells were passaged at 90% confluence using 0.05% Trypsin-EDTA and were routinely tested for mycoplasma contamination. For *vitro* experiments, the cells were seeded at density of ca.52000cells/cm^2^, 16-20hrs prior to stimulation with MG in DMEM with 0.1% FCS, 1 g/l glucose, 1% penicillin/streptomycin, 1% amphotericin B, 10mM HEPES at 37°C.

### Measurement of Methylglyoxal by LC-MS/MS

Methylglyoxal in cells and media was determined by isotope dilution, tandem mass spectroscopy, following derivatization with 1,2-diaminobenzene (Rabbani and Thornalley, 2014b). Briefly, cells (ca.1×10^6^ cells) were homogenized in ice-cold 20% (wt/vol) trichloroacetic acid in 0.9% (wt/vol) sodium chloride (20 μl) and water (80 μl). The media samples (ca.20 μl) were acidified ice-cold 20% (wt/vol) trichloroacetic acid in 0.9% (wt/vol) sodium chloride (10 μl) and water (20 μl). An aliquot (5 μl) of the internal standard (13C3-MG; 400 nM) was then added and the samples vortexed mixed. Following centrifugation (14000 rpm; 5 mins @ 4 °C), 35 μl of the supernatant was transferred to an HPLC vials containing a 200 μl glass interest. An aliquot (5 μl) of 3 % sodium azide (wt/vol) was then added to each sample followed by 10 μl of 0.5 mM DB in 200 mM HCl containing 0.5 mM diethylenetriaminepentaacetic acid (DETAPAC) in water. The samples were then incubated for 4hrs at room temperature, protected from the light. Samples were then analyzed by LC-MS/MS using an ACQUITY™ ultra-high-performance liquid chromatography system with a Xevo-TQS LC-MS/MS mass spectrometer (Waters, Manchester, UK). The column was a Waters BEH C18 (100 x 2.1 mm) and guard column (5 x 2.1 mm). The mobile phase was 0.1 % formic acid in water with a linear gradient of 0–100 % 0.1 % formic acid in 50 % Acetonitrile:water over 0-10mins; the flow rate was 0.2ml/min and column temperature was 5°C. The capillary voltage was 0.5 kV, the cone voltage 20 V, the interscan delay time 100 ms, the source and desolvation gas temperatures 150 and 350 °C, respectively, and the cone gas and desolvation gas flows were 150 and 800 l/h, respectively. Mass transitions (parent ion>fragment ion; collision energy), retention time, limit of detection, and recoveries were as follows: 145.0 > 77.1, 24eV, 5.93 min, 0.52 pmol and 98 %. Acquisition and quantification was completed with MassLynx 4.1 and TargetLynx 2.7 (Waters^®^).

### Measurement of D-lactate

D-lactate in cellular media was determined by enzymatic, endpoint, assay as previously described (McLellan et al., 1992).

### In-Cell Westerns for MG-H1

MCECs were seeded into a 96-well, black body, clear bottom plate at a density of 25000 cells per well in 200μl of 5% FCS DMEM media and allowed to adhere overnight. The media was removed and the cell monolayer washed once with PBS (200μl/well). 0.1% FCS DMEM media (100μl) is added to each well and the plate returned to the incubator for 1hr. Working solutions of MG were prepared in 0.1% FCS DMEM media immediately before at x2 the final concentration; 100μl of the dilution MG solution was then added to the appropriate wells to give a final concentration range 0-1000μM, and the plate returned to the incubator for 24hrs.The media was then discarded, the cell monolayer washed twice with PBS (200μl/well) and fixated with 4% PFA in PBS at room temperature for 15mins. The cells were then permeabilised with 0.25% Triton X100 in TBS (200μl/well) at room temperature for 15mins and then blocked with 10% goat serum in TBS (200μl/well) for one hour at room temperature. The cells were then incubated with MG-H1-x-biotin antibody (1:500 in 10% goat serum in TBS, 100μl per well) overnight at 4°C. The cells were then washed three times with 10% goat serum in TBS +0.1% Tween 20 (200μl/well), and then incubated with IRDye^®^ 800CW Streptavidin (LICOR;1:1000 in 10% goat serum in TBS + 0.1% Tween 20; 100μl/well) for one hour, at room temperature, protected from the light. The cells were then washed three times with 10% goat in TBS + 0.1% Tween 20 (200μl/well), twice with TBS + 0.1% Tween 20 (200μl/well). The cells were incubated with CellTag^™^ 700 Stain (LICOR; 1:500 in TBS) for one hour, at room temperature, protected from the light. The cells were then washed three times with TBS + 0.1% Tween 20 (200μl/well), twice with TBS (200μl/well). The plates were then scanned, dry, and analysed using an Odyssey^®^ DLx scanner using Odyssey^®^ imaging software. The MG-H1 signal was detected in the 800-channel (pseudo-green) and the cell tag reagent was detected in the 700-channel (pseudo-red). The intensity of MG-H1 channel was expressed as percentage of the intensity from the CellTag channel.

### Measurement of Glycation, Oxidation and Nitration by LC-MS/MS

Protein-bound MG-H1 in MCECs stimulated with MG was determined by isotope dilution, tandem mass spectroscopy, as previously described (AHMED and THORNALLEY, 2002; THORNALLEY et al., 2003; Rabbani et al., 2014). Briefly, total protein extracts from cells (ca.3×10^6^ cells) were prepared by homogenization in 10mM sodium phosphate buffer (pH 7.4). The soluble protein fraction was retained and then concentrated by microspin ultrafiltration (10 kDa cut-off) at 14000 rpm for 30 mins at 4°C, and then washed by five cycles of concentration. An aliquot of the washed protein (100μg/20μl) was then hydrolyzed by serial enzymatic digestion using pepsin, pronase E, aminopeptidase and prolidase as described previously. An aliquot of the sample (ca.30μl) was spiked with an equal volume of 0.2% TFA in water containing the isotopic standards (5-25 pmol). Normal and isotopic standards were either purchased (Cambridge Isotope, Polypeptide Laboratories, Iris Biotech) or prepared in-house, as described previously. Samples were then analyzed by LC-MS/MS using an ACQUITY ultra-high-performance liquid chromatography system with a Xevo-TQS LC-MS/MS spectrometer (Waters). Two 5μm HypercarbTM columns (Thermo Scientific) in series were used: 2.1 x 50mm, fitted with a 5 x 2.1mm pre-column, and 2.1 x 250mm. The mobile phases were 0.1% TFA in water and 0.1% TFA in 50% water. The column temperature and flow rate were 30°C and 0.2ml/min, respectively. Analytes were eluted using a two-step gradient and the columns washed after each sample with 0.1% TFA in 50% THF, as described previously (Rabbani et al., 2014; THORNALLEY et al., 2003)). MG-H1 and arginine were detected by electrospray positive ionization with multiple reaction monitoring (MRM). The ionization source temperature was 150°C and the desolvation temperature was 500°C. The cone gas and desolvation gas flows were 150 and 1000 L/hr, respectively. The capillary voltage was 0.5 kV. Molecular ion and fragment ion masses, as well as cone voltage and collision energizes were optimized to ± 0.1 Da and ± 1eV for MRM detection of the analytes. Acquisition and quantification was completed with MassLynx 4.1 and TargetLynx 2.7 (Waters^®^).

### Cell Viability Markers

MCECs were seeded into 96-well (i) clear body plate for total metabolic activity; (ii) 96-well, white body plate for ATP content; (iii) clear bottom, blacked body plate for DNA content or (iv) clear body plate for L-lactate dehydrogenase (LDH) release, and stimulated with MG for 24rs, as previously described. For total metabolic activity, MTT solution (2mg/ml in PBS) was then added to each well (50μl), covered in aluminium foil to protect from the light, and returned to the incubator for 3hrs. The media was removed and the DMSO (200μl) added to each well, to lysis the cells and solubilize the purple formazan crystal, and then incubated at room temperature on an orbital shaker for 1hr and the absorbance at 590nm (reference at 690 nm) measured. Total ATP content was measured using the ATPlite Luminescence Assay System (Perkin Elmer), according to the manufacturer’s instructions. Total DNA content was measured using the CyQUANT^™^ direct cell proliferation assay (Thermo), according to the manufacturer’s instructions and fluorescence intensity measured at Ex/Em 480/520nm. LDH release into the media was measured using the Pierce LDH cytotoxicity assay kit (Thermo), according to the manufacturer’s instructions and absorbance at 490nm (reference at 680nm) measured. LDH release was calculated as percentage of the maximum release, which was determined by addition of 10x lysis buffer to each well. All measurements were performed using FLUOstar OMEGA multiplate reader (BMG Labtech). For all markers, the respective data was calculated as a percentage of controls (untreated cells) and fitted by nonlinear regression to determine the EC_50_.

### Apoptosis & Cell Viability

MCECs were seeded into a 10cm petri dishes at a density of 3×10^6^ cells per well in 10ml of 5% FCS DMEM media and allowed to adhere overnight. The media was removed and the cell monolayer washed once with PBS (5ml/dish). 0.1% FCS DMEM media (6ml) was added to each dish and the dishes returned to the incubator for 1hr. Working solutions of MG were prepared in 0.1% FCS DMEM media immediately before at x2 the final concentration; 6ml of the dilution MG solution was then added to the appropriate wells to give a final concentration range 0-1000μM, and the dishes returned to the incubator for 24hrs. The media from each dish was collected, and the cells were harvested with Accutases and pooled with the appropriate media. The resulting cell suspension was then centrifuged (1500rpm; 5mins; 4°C), and the cell pellet resuspend in PBS (1ml) and split equally between two Eppendorf tubes (1.5ml). For the measurement of apoptosis, the cells were collected by centrifugation (1500rpm; 5mins; 4°C) and washed twice with Annexin V binding buffer (Biolegend; 1ml per wash). The cell pellets were resuspended in of binding buffer (100μl) and transferred to a well of 96-well ‘V’-shaped plate, and then centrifuged (2000rpm; 2mins; 4°C). The supernatant was discarded and the cell pellets were resuspended in binding buffer containing Alexafluor647-Annexin (Biolegend; 1:200 in binding buffer; 100μl per well) and incubated at room temperature, protected from the light, for 30mins. The plate was then centrifuged (2000rpm; 2mins; 4°C) and the cell pellets washed twice with binding buffer (100μl/well). The cell pellets were resuspended in 4% PFA (100μl/well) and incubated at room temperature, protected from the light, for 15mins. The plate was then centrifuged (2000rpm; 2mins; 4°C) and the cell pellets washed twice with binding buffer (100μl/well) and resuspend in PI solution (Sigma; 50μg/ml in binding buffer; 100μl/well) and stored at 4°C, protected from the light, until analyzed by flow cytometry. For the measurement of viability, the cells were collected by centrifugation (1500rpm; 5mins; 4°C) and washed twice with PBS (1ml per wash). The cell pellets were resuspended in PBS (100μl) and transferred to a well of 96-well ‘V’-shaped plate, and then centrifuged (2000rpm; 2mins; 4°C). The supernatant was discarded and the cell pellets were resuspended in PBS containing Zombie NIR^™^ dye (Biolegend; 1:200 in PBS; 100μl per well) and incubated at room temperature, protected from the light, for 30mins. The plate was then centrifuged (2000rpm; 2mins; 4°C), the cell pellets washed twice with PBS (100μl/well). The cell pellets were resuspended in 4% PFA (100μl/well) and incubated at room temperature, protected from the light, for 15mins. The plate was then centrifuged (2000rpm; 2mins; 4°C) and the cell pellets washed twice with 10% FCS in PBS (100μl/well). The cell pellets were then resuspended in a 10% FCS in PBS (100μl/well) and stored at 4°C, protected from the light. Flow cytometry analysis was performed using LSR II flow cytometer (Becton Dickinson Bioscience) equipped with three lasers (405 nm, 488 nm, and 633 nm) and a 96-well plate sample loader and the data analysed with FlowJo_V10.0.7 software (FlowJo LLC, Ashland, OR, USA). The cell populations were initially gated via forward scatter (FSC) against side scatter (SSC). For the apoptosis analysis, fluorescence distribution was displayed as a two-color dot plot analysis from which four different populations were identified: Dapi^−^/Annexin V^−^ (*viable cells*), Dapi^−^/Annexin V^+^ (*early apoptotic*), Dapi^+^/Annexin V^−^ (*dead*), Dapi^+^/Annexin V^+^ (*late apoptotic cells*). The frequency of fluorescent cells in each quadrant was determined. For the viability analysis, fluorescence distribution was displayed a histogram in which the frequency of Zombie^−^/Zombie^+^ cells were used as a measure on viability.

### Proliferation Analysis by Real-Time Cell Analysis (RTCA)

MCECs were seeded into E-plate, which is equivalent to the surface area of a 96-well plate and placed into a xCELLigence^®^ RTCA DP instrument, housed within a 37°C incubator. The adhesion of the cells to the electrode surface was monitored overnight. The media was removed and the cell monolayer washed once with PBS (200μl/well). 0.1% FCS DMEM media (100μl) was added to each well and the plate returned to the RTCA instrument for 1hr. Working solutions of MG were prepared in 0.1% FCS DMEM media immediately before at x2 the final concentration; 100μl of the dilution MG solution was then added to the appropriate wells to give a final concentration range 0-1000μM, and the plate returned to the incubator for proliferation measured every hour for a period of 48hrs.

### Cell Cycle Analysis, Quiescence & Senescence

MCECs were seeded into a 10cm petri dishes and stimulated with MG as previously described. After 24hrs, the media from each dish was collected, and the cells harvested with Accutases and pooled with the appropriate media. The resulting cell suspension was then centrifuged (1500rpm; 5mins; 4°C), and the cell pellet resuspend in PBS (1ml) and split equally between two Eppendorf tubes (1.5ml). For the measurement of the cell cycle and quiescence, cells were collected by centrifugation (1500rpm; 5mins; 4°C) and washed twice with PBS (1ml per wash) and then resuspend in ice-cold, 70% ethanol and incubated for one hour at −20°C. The cells were collected by centrifugation (1500rpm; 5mins; 4°C) and washed twice with 10% FCS in PBS (1ml per wash). The cell pellets were resuspended in 10% FCS in PBS (100μl) and transferred to a well of 96-well ‘V’-shaped plate, and then centrifuged (2000rpm; 2mins; 4°C). The supernatant was discarded and the cell pellets were resuspended in 10% FCS in PBS containing Alexafluor647 anti-Ki67 (Biolegend; 1:200 in binding buffer; 100μl per well) and incubated at room temperature, protected from the light, for 30mins. The plate was then centrifuged (2000rpm; 2mins; 4°C) and the cell pellets washed twice with 10% FCS in PBS and resuspend in Dapi solution (Sigma; 50μg/ml in 10% FCS in PBS; 100μl/well) and stored at 4°C, protected from the light, until analyzed by flow cytometry. For the measurement of senescence, the CellEvent^™^ senescence green flow cytometry detection kit (Thermo) was used, according the manufacturer’s instructions. Briefly, cells were collected by centrifugation (1500rpm; 5mins; 4°C) and washed twice with PBS (1ml per wash). The cell pellets were resuspended in PBS (100μl) and transferred to a well of 96-well ‘V’-shaped plate, and then centrifuged (2000rpm; 2mins; 4°C). The supernatant was discarded and the cell pellets were resuspended in PBS containing Zombie NIR^™^ dye (Biolegend; 1:200 in PBS; 100μl per well) and incubated at room temperature, protected from the light, for 30mins. The plate was then centrifuged (2000rpm; 2mins; 4°C), the cell pellets washed twice with PBS (100μl/well). The cell pellet was then fixated, washed in the assay buffer and resuspended in the working solution (100μl) and incubated for 2hrs at 37°C in a CO_2_-free incubator, protected from the light. The plate was then centrifuged (2000rpm; 2mins; 4°C) and the cell pellets washed twice with 10% FCS in PBS (100μl/well). The cell pellets were then resuspended in 10% FCS in PBS (100μl/well) and stored at 4°C, protected from the light. Flow cytometry was performed as previously described. For the analysis, the cell populations were initially gated via forward scatter against side scatter and data. Single cell gate was then created based upon FSC-H vs. FSC-A. For the analysis of cell cycle, the Dapi signal of the single cell gate was with acquired with linear amplification, and displayed as a histogram to visualize three phases of the cell cycle (G1, S and G2/M). The frequency of Ki-67 positive cells in in the G1 was then determined. For the analysis of senescence, the Zombie^−^ (viable) population was selected from single cell gate and mean GFP fluorescence intensity (β-galactosidase activity) determined.

### Macromolecule Synthesis by Click-iT Chemistry

MCECs were seeded into a 96-well, black body, clear bottom plate and stimulated with MG for 24rs, as previously described. For the measurement of DNA, RNA and protein synthesis, the cells were incubated for 2 hours with either 5-ethynyl-2’-deoxyuridine (EdU; 10μM) for DNA, 5-Ethynyl-uridine (5-EU; 1mM) for RNA, or L-Homopropargylglycine (L-HPG; 100μM) for protein. The media was then discarded, the cell monolayer washed twice with PBS (200μl/well) and fixated with 4% PFA in PBS at room temperature for 15mins. The cells were then permeabilised with 0.25% Triton X100 in TBS at room temperature for 15mins and then blocked with 1% BSA in TBS for one hour at room temperature. Click reaction was performed using the Click-iT^™^ cell reaction buffer kit (Thermo), according the manufacturer’s instructions, using IRDye^®^ 800CW Azide (LICOR). The plates were washed three times with TBS (200μl per well) and then scanned, dry, and analysed using an Odyssey^®^ DLx scanner using Odyssey^®^ imaging software. The synthesis of each macromolecule was expressed as a percentage of controls (untreated cells) and fitted by nonlinear regression to determine the EC_50_.

### Western Blot Analysis

Total protein extracts from MCECs stimulated with MG were prepared using RIPA buffer (50mM Tris-HCl (pH7.5), 150mM NaCl, 1% Triton X100, 0.5% Sodium deoxychloate, 0.1%SDS + 1mM DTT + protease/phosphatase inhibitor cocktail). Briefly, cell pellets were resuspended in RIPA buffer (300μl) and transferred to a bioruptor microtube (1.5ml) containing 100mg of protein extraction beads. The samples were sealed and then placed into a pre-cooled Bioruptor^®^ Pico and sonicated for 30 cycles of ON/OFF (Run time 30mins). The samples were then centrifuged (14000rpm; 5mins; 4°C) and the resulting supernatant was used for protein determination and further analysis. Cytoplasmic-nuclear fractionation from MCECs stimulated with MG were prepared using the NE-PER^™^ nuclear and cytoplasmic extraction kit (Thermo), according to the manufacturer’s instructions. Protein concentrations in the different cell extracts was performed using the Bradford assay (Bradford, 1976). Proteins extracts(ca.20μg) were resolved by SDS-PAGE (4 - 15 % Mini-PROTEAN TGX, Biorad) and transferred onto a nitrocellulose membrane and were blocked according to appropriate primary antibodies Membranes were visualized using an Odyssey^®^ DLx scanner using Odyssey^®^ imaging software. The primary and secondary antibodies used are as follows:

| Target | MW (kDa) | Company/ Catalogue No. | Isotype | Blocking/Diluent | Dilution |
| --- | --- | --- | --- | --- | --- |
| **Primary Antibodies** |  |  |  |  |  |
| PARP | 89/116 | CST/9532 | Rabbit IgG | 5% Milk in TBST | 1:1000 |
| Cleaved-PARP | 89 | CST/9544 | Rabbit IgG | 5% Milk in TBST | 1:1000 |
| Caspase-3 | 17/19/35 | CST/14220 | Rabbit IgG | 5% BSA in TBST | 1:1000 |
| Cleaved-Caspase-3 | 17/19 | CST/9664 | Rabbit IgG | 5% BSA in TBST | 1:1000 |
| β-Actin | 45 | CST/3700 | Mouse IgG2b | 5% BSA/Milk in TBST | 1:2000 |
| Phospho-p53 | 53 | CST/9284 | Rabbit IgG | 5% Milk in TBST | 1:1000 |
| Phospho-H2Ax | 15 | CST/9718S | Rabbit IgG | 5% BSA in TBST | 1:1000 |
| Cyclin B1 | 55 | CST/4138 | Rabbit IgG | 5% BSA in TBST | 1:1000 |
| Cyclin E1 | 48 | CST/20808S | Rabbit IgG | 5% BSA in TBST | 1:1000 |
| Phospho-CDK2 | 33 | CST/2561S | Rabbit IgG | 5% BSA in TBST | 1:1000 |
| Phospho-Histone H3 | 17 | CST/53348 | Rabbit IgG | 5% BSA in TBST | 1:1000 |
| **Secondary Antibodies** |  |  |  |  |  |
| Anti-rabbit IgG (H+L) (DyLight™ 800 4X PEG Conjugate) |  | CST/5151 | Goat IgG | 5% Milk/BSA in TBST | 1:10000 |
| Anti-mouse IgG (H+L) (DyLight™ 680 Conjugate) |  | CST/5470 | Goat IgG | 5% Milk/BSA in TBST | 1:10000 |

### Immunofluorescence Analysis

MCECs were seeded onto 12mm glass cover slips in 24-well and allowed to adhere overnight. The media was removed and the cell monolayer washed once with PBS (0.5ml/well). 0.1% FCS DMEM media (0.25ml) was added to each dish and the dishes returned to the incubator for 1hr. Working solutions of MG were prepared in 0.1% FCS DMEM media immediately before at x2 the final concentration; 0.25ml of the dilution MG solution was then added to the appropriate wells to give a final concentration range 0-1000μM, and the dishes returned to the incubator for 24hrs. Cells were fixated directly by addition of 4% PFA to each well, at final concentration of 0.2% and the incubated at 4°C for 16hrs. The media was discarded, and the cell monolayer washed three times with TBS. The cells were then permeabilised with 0.25% Triton X100 in TBS at room temperature for 15mins and then blocked with 5%BSA in TBS for one hour at room temperature. The cells were then incubated with anti-complex I antibody (Abcam, ab109798; 1:200 in 5% BSA in TBS, 200μl per well) overnight at 4°C. The cells were then washed three times with 5% BSA in TBS + 0.1% Tween 20, and then incubated with anti-mouse-alexafluor488-IgG (Cell signalling; 1:1000 in 5% BSA in TBS, 200μl per well) for one hour, at room temperature, protected from the light. The cells were then washed three times with 5% BSA in TBS + 0.1% Tween 20, once with TBS and the incubated with Phalloidin-iFluor 647 (Abcam; 1:200 in TBS; 200μl per well) at room temperature for 30mins, protected from the light. The cells were then washed three times with 5% BSA in TBS + 0.1% Tween 20, once with TBS, stained with Dapi and mounted with Permaflour mountant medium (Thermo). Images were acquired on Zeiss LSM780 microscope with oil immersion at x40 magnification and analyzed using ImageJ software.

For the visualization of MG-H1, a monoclonal rat Ig2C antibody (clone 6D7) was used (Morgenstern et al., 2017). The antibody was conjugated to biotin using the DSB-X^™^ biotin protein labelling kit (Thermo), according to the manufacturer’s instructions. In order to use this antibody (1:500 dilution), the following modifications were made to the protocol: (*i*) BSA was replaced with goat serum, to reduce non-specific binding and (*ii*) avidin/biotin blocking was performed prior to addition of the MG-H1 antibody using the Avidin/Biotin Blocking Kit (Vector Laboratories), according to the manufacturer’s instructions. The MG-H1 signal was visualized using Alexafluor488-Streptavidin (Biolegend; 1:500).

### Characterization of Endothelial Activation Status

MCECs were seeded into a 6-well plates in 2ml of 5% FCS DMEM media and allowed to adhere overnight. The media was removed and the cell monolayer washed once with PBS (2ml/dish). 0.1% FCS DMEM media (1ml) was added to each well and the plates returned to the incubator for 1hr. Working solutions of MG were prepared in 0.1% FCS DMEM media immediately before at x2 the final concentration; 1ml of the dilution MG solution was then added to the appropriate wells to give a final concentration range 0-500μM, and the dishes returned to the incubator for 24hrs. The cells were harvested with Accutases, collected by centrifugation (1500rpm; 5mins; 4°C), and washed twice with PBS (1ml per wash). The cell pellets were resuspended in PBS (100μl) and transferred to a well of 96-well ‘V’-shaped plate, and then centrifuged (2000rpm; 2mins; 4°C). The supernatant was discarded and the cell pellets were resuspended in PBS containing Zombie Violet dye (Biolegend; 1:200 in PBS; 100μl per well) and incubated at room temperature, protected from the light, for 30mins. The plate was then centrifuged (2000rpm; 2mins; 4°C), the cell pellets washed twice with 10% FCS in PBS (100μl/well) and then resuspended in 10% FCS in PBS (100μl/well) containing TruStain FcX^™^ (anti-mouse CD16/32) antibody (Biolegend; 1:200) and incubated on ice for 20mins, protected from the light. The plate was then centrifuged (2000rpm; 2mins; 4°C) and the cell pellets resuspended in 10% FCS in PBS (100μl/well) containing AlexaFluor488-CD106/VCAM-1, PE/Dazzle594-CD54/ICAM-1, CD62E/E-selectin, PerCP/Cyanine5.5-CD146/MUC18, and APC-CD201/EPCR (Biolegend; 1:250) and incubated on ice for one hour, protected from the light. The plate was then centrifuged (2000rpm; 2mins; 4°C), the cell pellets washed twice with 10% FCS in PBS (100μl/well) and then resuspended in 10% FCS in PBS (100μl/well) containing APC/Cyanine7 Goat anti-mouse IgG (Biolegend; 1:250) and incubated on ice for 30mins, protected from the light. The plate was then centrifuged (2000rpm; 2mins; 4°C), the cell pellets washed twice with 10% FCS in PBS (100μl/well) and then resuspended in 4% PFA (100μl/well) and incubated at room temperature, protected from the light, for 15mins. The plate was then centrifuged (2000rpm; 2mins; 4°C) and the cell pellets washed twice with 10% FCS in PBS (100μl/well). The cell pellets were then resuspended in a 10% FCS in PBS (100μl/well) and stored at 4°C, protected from the light. Flow cytometry was performed as previously described. For the analysis, the cell populations were initially gated via forward scatter against side scatter and data. Single cell gate was then created based upon FSC-H vs. FSC-A. Compensation analysis between overlapping fluorophores was performed using single-antibody stained samples, generated from a mixed population of cells. From the single cell gate, the Zombie-(viable) population was selected and the mean fluorescence intensity for each marker determined.

### Endothelial Adhesion Assay

MCECs were seeded into 6-well plates and stimulated with MG as previously described. After 24hrs the media was discard, the cell monolayer was washed once with PBS and 0.1% FCS DMEM media (1ml) added to each well and returned to the incubator. The mouse macrophate cell line, J774A.1 (cultured in 10% FCS, 1% P/S, in high-glucose, DMEM) were harvested, counted and labelled with carboxyfluorescein succinimidyl ester (CFSE) (Biolegend; 5μM in PBS). The CFSE-labelled J774A.1 cells were collected by centrifugation (1500rpm; 5mins; 4°C), washed three times with growth media and resuspend at a density of 2×10^6^ cells/ml. 1ml of the CFSE-labelled J774A.1 cells was added to each well of the 6-well plate and incubated for one hour at 36°C. The media was subsequently discarded and the cell monolayer washed five times with PBS + 0.2mM CaCl_2_, 0.1mM MgCl_2_. The cells were then harvested with Accutases, collected by centrifugation (1500rpm; 5mins; 4°C), and washed twice with PBS (1ml per wash). The cells were then resuspended in 10% FCS in PBS (100μl) and transferred to a well of 96-well ‘V’-shaped plate, and stored at 4°C, protected from the light. Flow cytometry was performed as previously described. For the analysis, the cell populations were initially gated via forward scatter against side scatter and data and the number of CFSE+ expressed as a percentage of the total population.

### TNFα Responsiveness of Endothelial Cells

MCECs were seeded into a 6-well plates and stimulated with MG as previously described. After 24hrs, the cells were stimulated with TNFα (10ng/ml; Biolegend) and incubated for 16hrs at 37°C. The media from each dish (ca.1ml) was centrifuged (14000rpm; 5mins; 4°C) to pellet cell debris and then transferred to an appropriate labelled Eppendorf (1.5ml) and stored at −20°C. The cells were harvested, collected, proceed and analysed as previously described for the assessment of the activation status. A second set of 6-well plates were setup in parallel for isolation of total RNA and qPCR analysis.

### Wound Healing

MCECs were seeded Incucyte^®^ Imagelock 96-well plates and stimulated with MG as previously described. After 24hrs the media was exchanged and a linear scratch was made, using the Incucyte ^®^ Wound Maker. The progression of wound healing was then monitored every hour for a total of 72hrs using the IncuCyte ZOOM ^®^. Relative wound density was calculated using the IncuCyte ^™^ Chemotaxis Cell Migration Software.

### Angiogenesis Assay

Tube forming assay was performed as previously described (Arnaoutova and Kleinman, 2010). Briefly, MCECs were seeded into a 10cm petri dishes and stimulated with MG as previously described. After 24hrs, the cells were harvested with Accutases, collected by centrifugation (1500rpm; 5mins; 4°C), washed twice with PBS (1ml per wash) and labelled with CFSE, as previously described. The CFSE-labelled MCECs were resuspended at a density of 1.2×10^6^ cells/ml and aliquot (100μl) added to the well of 96-well, black body, clear bottom plate containing solidified matrigel (50μl). The plate was then incubated at 37°C for four hours, after which the extent of tube formation was assessed by fluorescence microscopy using Zeiss Axio Vert. A1 microscope at x5 magnification. Quantitative analysis of angiogenesis was performed using ImageJ Angiogenic Analyzer

### General Adhesion Assay

MCECs were seeded into a 10cm petri dishes and stimulated with MG as previously described. After 24hrs, the cells were harvested with Accutases, collected by centrifugation (1500rpm; 5mins; 4°C), washed twice with PBS (1ml per wash) and labelled with CFSE, as previously described. The CFSE-labelled MCECs were resuspended at a density of 0.25×10^6^ cells/ml and aliquot (100μl) added to the well of 96-well, black body, clear bottom plate. The plate was then incubated at 37°C for six hours. The media was discarded and the cells washed three times with PBS (200μl/well) and fluorescence intensity at Ex/Em 480/520nm was measured using FLUOstar OMEGA multiplate reader (BMG Labtech) and expressed as percentage of the controls (untreated cells).

### Endothalial Barrier Function

MCECs (50,000 cells) were seeded onto gelatin coated on a polyester/polycarbonate mesh was seeded and cultured on a polyester/polycarbonate mesh (Transwell, 0.4 μm pore size, 12-well type; Costar, MA, USA). The apical and basolateral chambers of the Transwell were filled with 0.2ml and 1ml culture medium, respectively. Transendothelial electrical resistance (TER) was measured daily using an EVOM volt ohm meter equipped with STX-2 electrodes (World Precision Instruments, Sarasota, FL, USA) and electrical resistance calculated as previously described (Bartosova et al., 2021). Once a stable monolayer had formed (ca.2 days; TER was >10 Ω·cm^2^), cells were stimulated with MG, as previously described. After 24hrs, the TER was then measured. The paracellular permeability of the treated monolayers was determined by measuring the flux of 4 kDa fluorescein isothiocyanate (FITC) labelled dextran (obtained from Sigma Aldrich, Taufkirchen, Germany) from the apical to the basolateral compartment of a Transwell chamber. Once the TER measurements had been performed, 1mg/ml FITC-dextran was added to apical compartment of a Transwell chamber and the increase of the fluorescence intensity in the basolateral Transwell compartment measured after 4hrs. An equimolar amount of unlabeled dextran was added to the basolateral compartment of the Transwell system to maintain an isotonic condition. After 4hrs, an aliquot of media (ca.50μl) was taken from the basolateral compartment and the fluorescence intensity at Ex/Em 480/520nm measured using FLUOstar OMEGA multiplate reader (BMG Labtech). A calibration curve of FITC-dextan was used to calculation of amount which had been transported in the basolateral compartment and the results were expressed relative to Transwells which had not been seeded with cells. The transwells were then fixated and stained with Alexafluor555-ZO-1 an DAPI, as previously described (Bartosova et al., 2021). Images were acquired by fluorescence microscopy (ACQUIFER Imaging GmbH, Heidelberg, Germany) with equal exposure times and grey scale images analysed using ImageJ software.

### Quantitative PCR (qPCR)

Total RNA from MG stimulated MCECs was extracted using peqGOLD MicroSpin Total RNA kit (Peqlab) and then converted into cDNA with a high capacity cDNA reverse transcription kit (Thermo Fisher Scientific), according to the manufacturer’s instructions. qPCR was performed using DyNAmo ColorFlash SYBR Green qPCR Master Mix (Thermo Fisher Scientific) and a LightCycler 480 Instrument II (Roche Applied Science). Signals of amplified products were verified using melting curve analysis, and mRNA levels were normalized to the geometric mean of GAPDH and ß-actin. Relative expression levels were calculated using the Ct method (Livak and Schmittgen, 2001). Primer sequences are as follows:

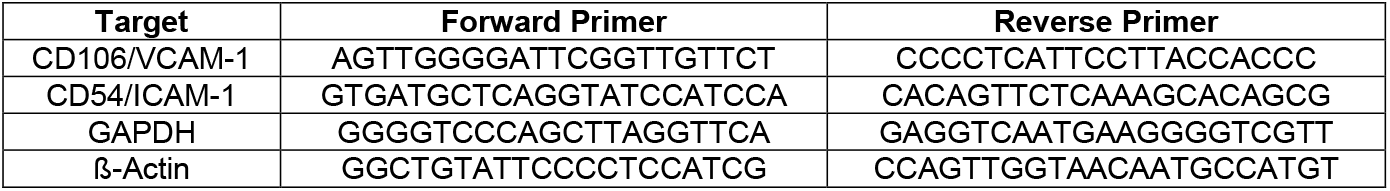

### Enzyme-linked immunosorbent assay (ELISA)

VCAM-1 in the media (1:100) from MCECs stimulated with MG was measured using commercial kits from Sigma, according to the manufacturer’s instructions.

### Proteases & Proteome Activity Assay

MCECs were seeded into a 96-well, white body, plates and stimulated with MG for 24rs, as previously described. Chymotrypsin, trypsin and caspase activity was measuring using the Cell-Based Proteasome-Glo^™^ Assays (Promega) accordingly to the manufacturer’s instructions. The inhibition of each proteases was expressed as a percentage of controls (untreated cells) and fitted by nonlinear regression to determine the EC_50_. Total proteasome activity was measured in MCECs stabling proteasome-sensitive fluorescent reporter, ZsProSensor-1 (Takara Bio, Tokyo, Japan). This reporter consists of a green fluorescent protein, ZsGreen, fused to mouse ornithine decarboxylase, which is degraded by the proteasome without being ubiquitinated. Under steady-state conditions, the fluorescent reporter is rapidly degradation. However, when the proteasome is inhibited, it is stabilized and the fluorescence intensity increases. Briefly, MCECs (1×10^6^ cells) were transfected 10μg ZsProSensor-1 plasmid using a NEON^®^ electroporation transfection system (Thermo) with the following conditions: pulse voltage: 1200mV; pulse width: 30ms; pulse number 1. G418-resistant, GFP-positive clones were selected by single-cell sorting using FACSAriaII (Becton Dickinson Bioscience) and expanded. Subsequent clones were screened with respect to expression, by immunofluorescence, and functionality, by flow cytometry, following stimulation with the proteasome inhibitor, MG132. Those clones which were positive for both aspects were subsequently used. MCECs were seeded into a 96-well, black body, clear bottom plate and stimulated with MG for 24rs, as previously described.The plate was then washed three times with PBS (200μl per well) and the fluorescence intensity measured at Ex/Em 480/520nm (ZsProSensor-1/GFP) and 360/460nm (Nuclei/DAPI) using CLARIOstar Plus (BMG Labtech). The GFP-to-DAPI was then determined and the inhibition of the proteasome expressed as a percentage of controls (untreated cells) and fitted by nonlinear regression to determine the EC_50_.

### Electron Microscopy

MCECs were seeded onto 12mm glass cover slips in 24-well and stimulated with 500μM MG for 24-96hrs, or treated with proteasome inhibitor, MG132. The media was removed and the cells were fixated with 3% glutaraldehyde in 0.1 M cacodylate buffer (pH 7.4) for 90mins. The cells were then washed with 0.1 M cacodylate buffer, post-fixated for 30 min in 1% osmium tetroxide and again washed with 0.1 M cacodylate buffer. The cells were then dehydrated by a graded series ethanol, dipping in propylene oxide and embedded in Epon Araldite. The coverslips were then removed with heat and razor blades. Semithin and 70–80-nm ultrathin sections were cut with a Reichert Ultracut E ultramicrotome, contrasted with uranyl acetate and lead citrate, and examined with a JEOL JEM 1400 electron microscope. Images were acquired using a TVIPS F216 digital camera and analyzed using ImageJ software.

### Statistical analysis

GraphPad Prism version 9.2.0 and Microsoft Excel were used for analysis. Data represents mean ± SD, unless otherwise stated, and analysed using two-tailed unpaired t-test with Welch’s correction. The comparison of more than one group was achieved using an ordinary one-way or two-way ANOVA analysis followed by comparing all groups using Tukey’s (one-way ANOVA) or Sidak’s (two-way ANOVA) multiple comparison test. Linear regression and Pearson correlation coefficient was used to study associations between measured variables. Differences were considered significant at p < 0.05

## Supplementary Figure Legends

**Supplementary Figure 1.** Wound healing capacity of MCECs stimulated with MG (0-500μM) was measured using the Incucyte^®^ System, as described in material and methods section. **(A**) Representative phase-contrast images of the wound closure for the MCECs stimulated with different concentrations of MG at 24 and 48hrs. Relative wound density was calculated using the IncuCyte ^™^ Chemotaxis Cell Migration Software (**B**). The dotted lines the time points at which the representative images shown in the upper panel were taken. (**C**)The rate of wound closure (%/hr) was calculated from the relative wound density curves by fitting to an exponential plateau using GraphPad Prism^®^ software (*right panel*). Data shown as min-to-max box plot with the median indicated by the line in the middle of the box (N=8); ****P ≤ 0.0001, **P ≤ 0.01 vs. unstimulated MCECs. Angiogenetic potential of MCECs stimulated with MG was measuring by Matrigel-based tube-forming assay, as described in the material and methods section. Images were acquired on Zeiss Axio Vert. A1 microscope at x5 magnification. (**D**) Representative images of the tube formation for the MCECs stimulated with different concentrations of MG. Scale bar represents 400μm. Quantitative analysis of angiogenesis was performed using ImageJ Angiogenic Analyzer (**E**). Data represents mean ± SD (N = 4); ****P≤ 0.0001, ***P ≤ 0.001, **P ≤ 0.01, *P ≤ 0.05 vs. unstimulated MCECs. Barrier function of MCECs stimulated with MG was measuring by transendothelial electrical resistance (TER; **F**) and FITC-dextran leakage (**G**), 24hr post-stimulation, as described in the material and methods section. Data represents mean ± SD (N = 3); **P ≤ 0.01 vs. unstimulated MCECs. MCECs were also fixated, permeabilized, blocked and stained for ZO1 (Red), and nuclei (Blue; Dapi), as described in the material and methods. Images were acquired on an ACQUIFER Imaging system at x20 magnification. (**H**) Representative images of ZO1 staining in the MCECs stimulated with different concentrations of MG. Scale bar represents 20μm. The intensity of ZO1 at the membrane was quantified using ImageJ and expressed as function of the area of interest (**I**). Data represents mean ± SD (N = 3); ****P ≤ 0.0001, ***P ≤ 0.001, *P ≤ 0.05 vs. unstimulated MCECs.

**Supplementary Figure 2.** The effect of media exchange on DNA content and synthesis of DNA, RNA and protein (**A**). For comparison, the MG-H1 data from Figure 4 is included. Data represent mean ± SD (N=8 per time point); ****P ≤ 0.0001, *P ≤ 0.01 vs. 24hr time point. (**B**) The inhibition of DNA, RNA and protein synthesis in MCECs stimulated with MG after 24hrs was measured as described in the material and methods. Data represents mean ± SD (N = 8) and EC_50_ curves generated using GraphPad Prism^®^ software. (**C**) The inhibition of the proteasome in MCECs stimulated with MG after 24hrs was measured as described in the material and methods. Data represents mean ± SD (N = 8) and EC_50_ curves generated using GraphPad Prism^®^ software. (**D**) Electron microscopy analysis of MCECs stimulated with and without MG, 72hrs-post-stimulation (scale bar 3μm), showing no morphological differences and MCECs treated with the proteasome inhibitor, MG132, for 24hrs (scale bar 500μm). Encircled area indicates electrodense deposits resulting from impaired protein degradation.

**Supplementary Figure 3.** MG-H1 staining of normal, cardiovascular disease (CVD) and tumor derived cardiovascular tissues. Tissue microarray (TMA) consisting a variety of cardiovascular tissues was stained for MG-H1 (Red) and nuclei (Blue; Dapi): Images were acquired on a Zeiss LSM780 confocal microscope with a Plan-Apochromat 20x objective. Representative images are shown in the *upper* panel in which the scale bar represents 100μm. Quantification of the total MG-H1 signal in cardiovascular tissues (*lower* panel, *left* side). Data is shown as min-to-max box plots, with the median indicated by the line in the middle of the box (N=7-9). Frequency of MG-H1 Positive Nuclei in cardiovascular tissues (*lower* panel, *middle*). Data is shown as min-to-max box plots, with the median indicated by the line in the middle of the box (N=7-9); **P ≤ 0.01, *P ≤ 0.05 vs. normal tissue. Frequency of MG-H1 Positive Nuclei in cardiovascular tissues accordingly to diagnosis (*lower* panel, *right* side). Data represent mean ± SD (N=2); ****P ≤ 0.0001, **P ≤ 0.01, *P ≤ 0.05 vs. normal tissue.

## Author contributions

**T.F., P.P.N** conceived, designed and supervised the study. **T.F., B.v.N., J.M., M.C., M.L.M, M.B.** performed experiments, collected analyzed the data. **J.M.** provided support with the MS analysis. **C.S.** provided the assistance with the assessment of human tissue. **I.H**. performed the EM analysis. **A.F**. provided support with respect to the characterization of the endothelial cells. **J.M., J.S.**, and **S.H.** provided support in conceiving and writing of the manuscript.

All authors approved the final version of the manuscript to be published

## Acknowledgments

The authors would like to thank Elisabeth Kliemank and Ulrike Ganserer for technical assistance as well members of the Nawroth/Fleming laboratory for helpful comments and discussions. The authors would also like to thank Dr. Holger Lorenz & Christian Hoerth, at the ZMBH Imaging facility, Dr. Walter Mier and his team in the Department of Nuclear Medicine, University Hospital of Heidelberg, Heidelberg, Germany and Dr. Tobias Dick and his team at Division of Redox Regulation, German Cancer Research Center (DKFZ), Heidelberg, Germany.

