## Supplementary figures and images for "Methylglyoxal Induces Endothelial Dysfunction via Stunning-like Phenotype"

### Supplmentary Figure 1

**Supplementary Figure 1.**

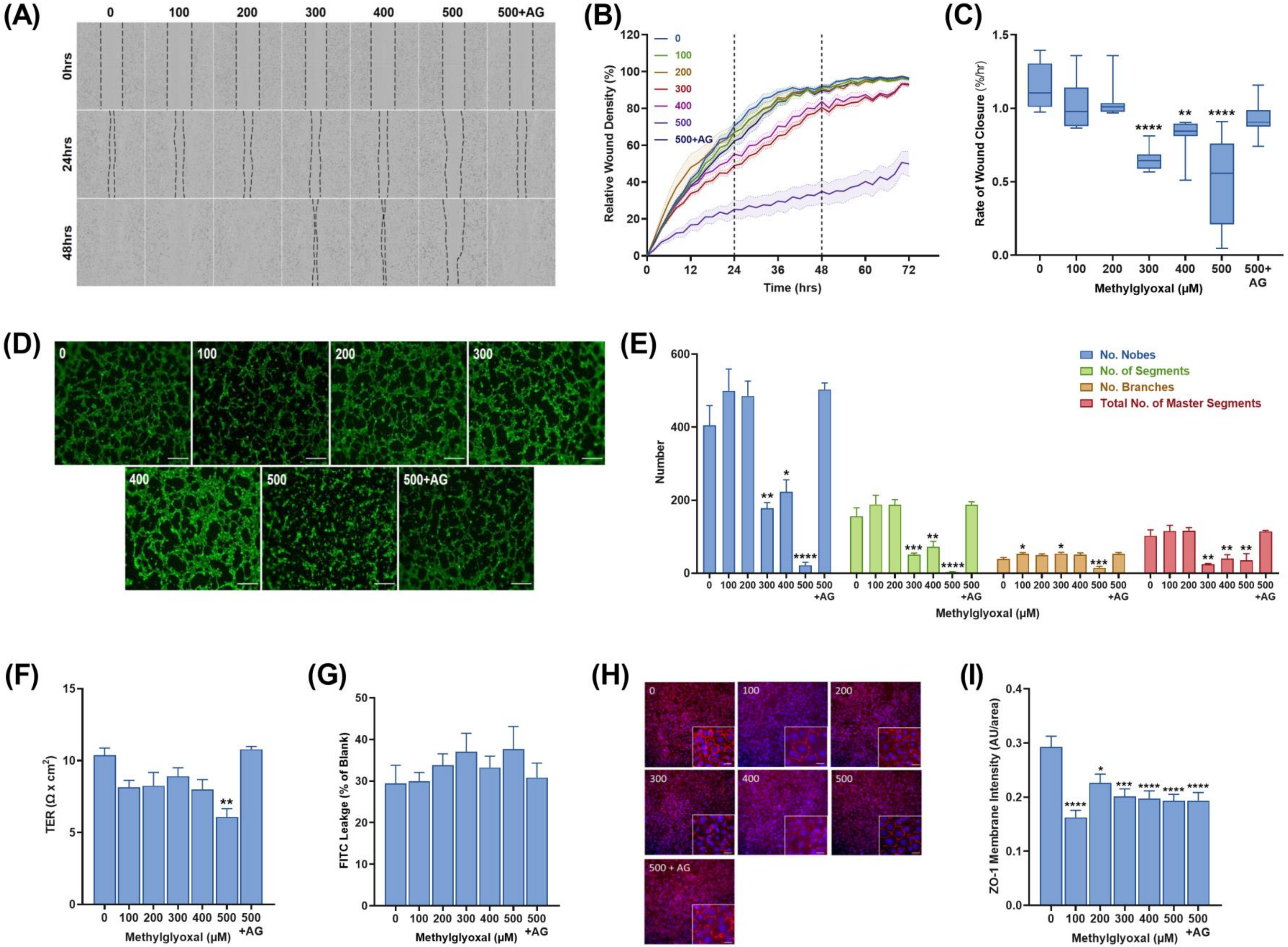

### Supplmentary Figure 2

#### Supplementary Figure 2.

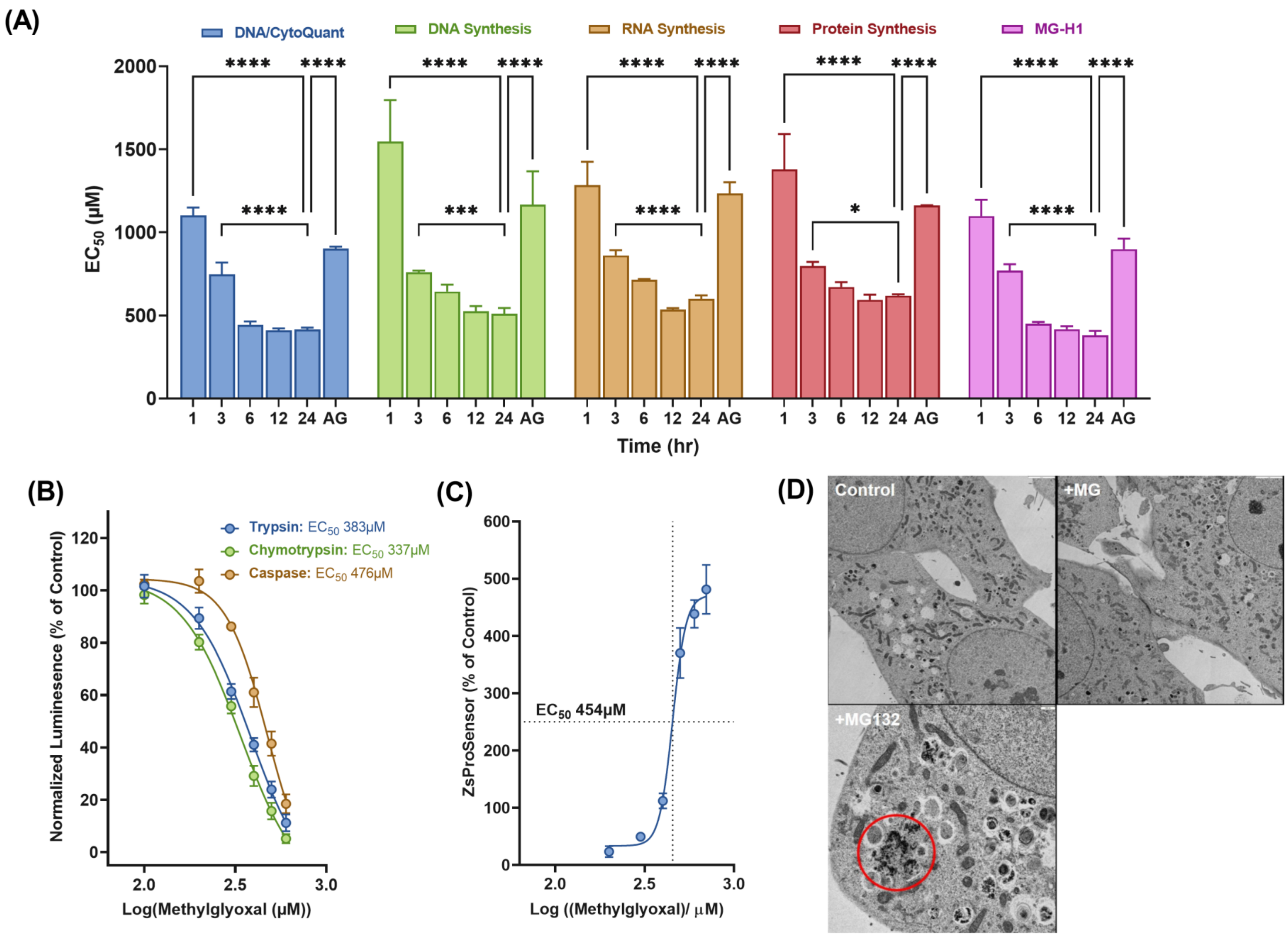

### Supplmentary Figure 3

**Supplementary Figure 3.**

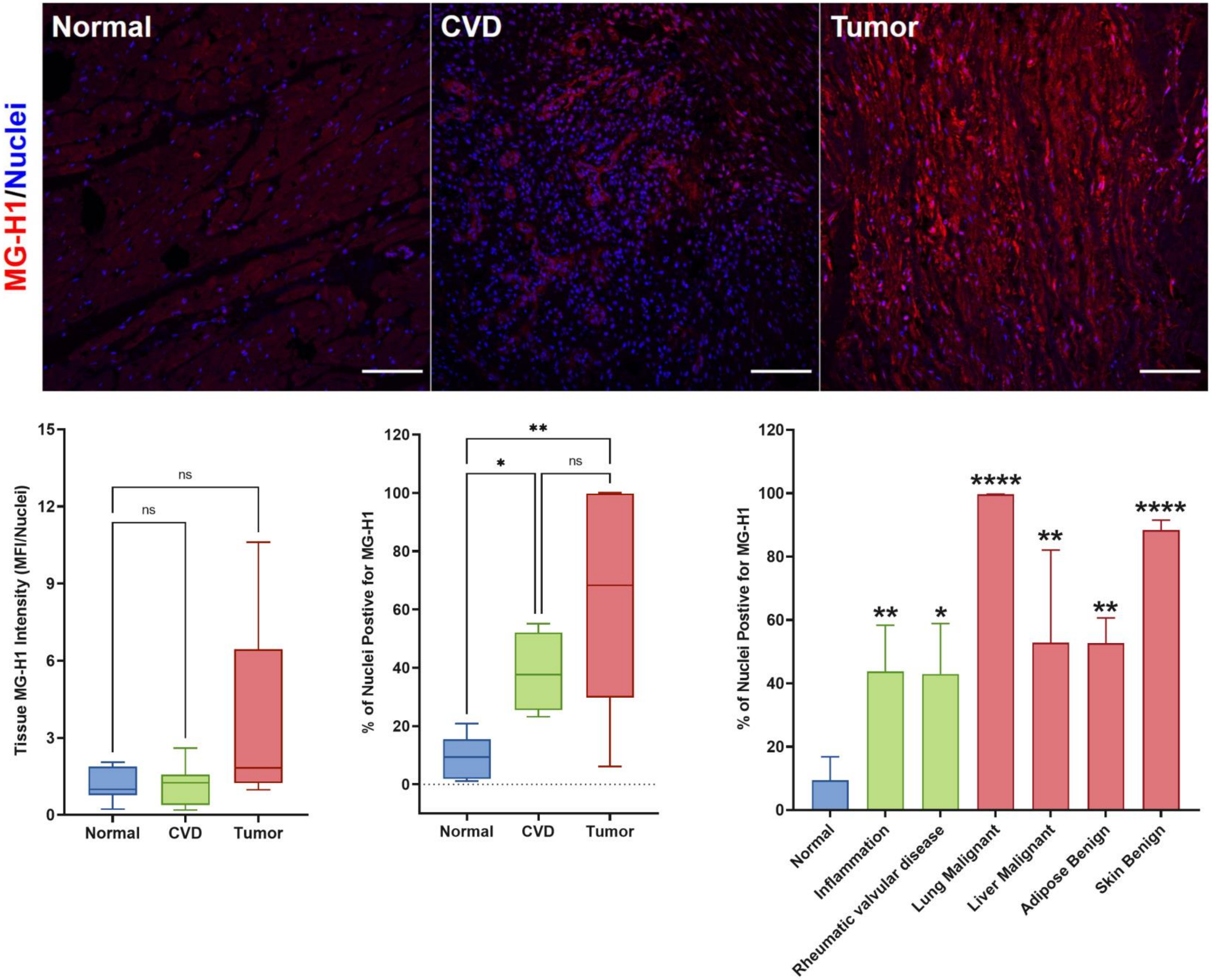
